# Small-molecule Polθ inhibitors provide safe and effective tumor radiosensitization in preclinical models

**DOI:** 10.1101/2022.09.27.509658

**Authors:** Gonzalo Rodriguez-Berriguete, Marco Ranzani, Remko Prevo, Rathi Puliyadi, Nicole Machado, Hannah R. Bolland, Val Millar, Daniel Ebner, Marie Boursier, Aurora Cerutti, Alessandro Cicconi, Alessandro Galbiati, Diego Grande, Vera Grinkevich, Jayesh Majithiya, Desiree Piscitello, Eeson Rajendra, Martin Stockley, Simon J. Boulton, Ester M. Hammond, Robert Heald, Graeme C. M. Smith, Helen Robinson, Geoff S. Higgins

**Affiliations:** Department of Oncology, University of Oxford; Old Road Campus Research Building, Roosevelt Drive, Oxford OX3 7DQ, United Kingdom; Artios Pharma; Babraham Research Campus, Cambridge, CB22 3FH, United Kingdom; Target Discovery Institute, Nuffield Department of Medicine, University of Oxford; NDM Research Building, Old Road Campus, Oxford, OX3 7FZ, United Kingdom; The Francis Crick Institute; London, NW1 1AT, United Kingdom

## Abstract

DNA polymerase theta (Polθ) is a DNA repair enzyme critical for microhomology mediated end joining (MMEJ). Polθ has limited expression in normal tissues but is frequently overexpressed in cancer cells and, therefore, represents an ideal target for tumor-specific radiosensitization. Here, we show that ART558 and ART899, two novel and specific allosteric inhibitors of the Polθ DNA polymerase domain, potently radiosensitize tumor cells, particularly when combined with fractionated radiation. Importantly, normal fibroblasts are not radiosensitized by Polθ inhibition. Mechanistically, we show that the radiosensitization caused by Polθ inhibition is most effective in replicating cells and is due to impaired DNA damage repair. We also show that radiosensitization is still effective under hypoxia, suggesting that these inhibitors may help overcome hypoxia-induced radioresistance. In addition, we describe for the first time ART899 and characterize it as a potent and specific Polθ inhibitor with improved metabolic stability. *In vivo*, the combination of Polθ inhibition using ART899 with fractionated radiation is well tolerated and results in a significant reduction in tumor growth compared to radiation alone. These results pave the way for future clinical trials of Polθ inhibitors in combination with radiotherapy.

## INTRODUCTION

Approximately 50% of cancer patients receive radiotherapy yet, despite technical improvements in delivery, survival rates following radical radiotherapy remain poor for many tumor types (1,2). Additionally, the radiation delivered to adjacent tissues is still frequently associated with significant side-effects. One strategy to improve radiotherapy outcome is to increase the radiosensitivity of tumor cells without affecting the sensitivity of the surrounding normal cells (3). DNA Polymerase theta (Polθ), a DNA repair enzyme that has low or absent expression in most normal tissues, but which is frequently overexpressed in many cancer types, represents an ideal tumor-specific radiosensitization target (4-8).

Polθ, encoded by the POLQ gene, plays a key role in microhomology-mediated end-joining (MMEJ) (9-12), a DNA double-strand break (DSB) repair pathway that depends on the presence of short homologous sequences (2-4 bp) across the break site. MMEJ has often been described as a back-up pathway for non-homologous end-joining (NHEJ) or repair by homologous recombination (HR) (13), but in specific circumstances MMEJ can function when both NHEJ and HR are active (14-16). Similarly to HR, MMEJ requires 5’-3’ resection of the DNA break, but during MMEJ the resection is much shorter and serves to reveal small microhomologies between the two single-stranded DNA (ssDNA) strands to enable their annealing (17,18). In contrast to HR –which is broadly considered error-free– MMEJ is an error-prone DSB repair pathway associated with deletions and introduction of single-base errors (19).

Polθ deficiency is synthetically lethal with defects in HR repair, suggesting that cancer cells with deficiency in HR, for example through BRCA mutations, become more reliant on the MMEJ pathway to repair DSBs and potentially other lesions (7,20). However, we have also shown that depleting Polθ through siRNA radiosensitizes tumor cells without mutations in BRCA genes, which suggests that Polθ inhibition may provide radiosensitization for a wide range of tumors, irrespective of HR status (21). Therefore, we initiated a Polθ inhibitor discovery program and recently described the identification of novel and specific Polθ inhibitors, including ART558 (22). These compounds, which specifically inhibit the polymerase activity of Polθ, were shown to be synthetically lethal with BRCA and Shieldin deficiency (22).

Here, we show that small molecule inhibitors targeting Polθ effectively radiosensitize tumor cells *in vitro* and *in vivo* and reveal their effect under hypoxia, a feature frequently associated with solid tumors and linked to radioresistance. Our results evidence that the radiosensitization induced by Polθ inhibition is cell cycle-dependent and caused by defects in DSB repair and increased genomic instability. We also report for first time ART899 as a specific and potent Polθ inhibitor with improved stability. Combined with fractionated radiation, Polθ inhibition exhibits effective tumor growth delay in mouse xenografts and, importantly, this combination treatment is well tolerated.

## METHODS

### Cell culture and reagents

H460, HCT116, T24, HeLa and MRC-5 cells were purchased from ATCC; AG01552 cells were obtained from the Coriell Institute. Cells were authenticated by short tandem repeat (STR) profiling (carried out by LGC standards) if grown for more than six months accumulatively after acquisition. U2OS Polθ CRISPR KO cells were generated by Synthego, which also provided the U2OS WT cells. Cells were grown in RPMI (H460, T24), DMEM (HCT116, HeLa), MEM (MRC-5, AG01552) or McCoy’s medium (U2OS), all supplemented with 10% FBS and incubated at 37°C and 5% CO_2_. All medium was purchased from Sigma-Aldrich/Merck; FBS was purchased from Life Technologies and the same serum batch was used for all experiments. Regular testing with MycoAlert kit (Lonza) confirmed the absence of mycoplasma contamination. ART558 was produced as described previously (22). ART899, was produced as described for its non-deuterated form, ART812 (22), with the addition of deuterium during synthesis. Compounds were stored as powder under vacuum at room temperature in the dark. Compounds were dissolved in DMSO at 12 mM and these stocks dissolved were kept at room temperature in the dark.

### Clonogenic survival experiments

Cells were plated as single cells in 6-well plates or 24-well plates and left to settle for a minimum of 5 hours before treatment. Seeding densities were optimized for each cell line; increasing cell numbers were used for higher IR doses to account for cell death. One hour prior to IR, a concentrated working stock of ART558 or ART899 was prepared in medium and added to the cells to yield a final concentration of 1 µM or 3 µM. Cells were irradiated in a cesium-137 irradiator (GSR D1 from Gamma Service; dose rate 1.2 Gy/min). Compound was removed by performing a medium change 3 days after IR. Colonies were grown for 8-14 days, stained with crystal violet and counted using the Gelcount automated colony counter (Oxford Optronics). The plating efficiency (PE = average colony number/cells plated) and the surviving fraction (SF = PE_IR Dose_/PE_0 Gy_) at a given IR dose was calculated. Survival data was fitted according to a linear quadratic equation and, for comparison between curves, the sensitization enhancement ratio at a surviving fraction of 0.10 (SER_10_) was calculated. The oxygen enhancement ratio (OER) was defined as the ratio between the radiation doses at 10% survival of hypoxic and normoxic cells.

### Hypoxia experiments

Hypoxia experiments were performed in a BactronEZ anaerobic chamber (Shel Lab) for concentrations of <0.1% O_2_ or a M35 hypoxia workstation (Don Whitley Ltd) for concentrations of 0.5% O_2_. Both chambers were humidified and kept at 37°C. Cells were left to settle at 37°C in a standard (normoxic) 37°C incubator for a minimum of 6 hours before transfer into the hypoxia chamber and were then left inside the chamber for at least 14 hours prior to IR. To perform radiations under hypoxic conditions, culture plates were transferred into airtight Perspex boxes that had also been placed inside the hypoxia chamber overnight (23). The boxes containing the culture plates were then placed inside the cesium irradiator for irradiation, thus maintaining hypoxia during irradiation. After radiation, culture plates were removed from the Perspex boxes and returned to a normoxic 37°C incubator for colonies to grow.

### Western blotting

Cells were washed in PBS and lyzed in RIPA buffer (ThermoFisher) supplemented with protease inhibitors (Roche) and benzonase (Merck). Lysates were cleared by centrifugation and protein content was quantitated using the Bicinchoninic acid assay (BCA; ThermoFisher). Equal amounts of samples (50-100 µg) were resolved by SDS-polyacrylamide gel electrophoresis in 3–8% Tris-Acetate gels (Thermofisher). Gels were wet-transferred in 1x NuPAGE Transfer Buffer (Thermofisher) 20% Ethanol and 0.05% SDS to nitrocellulose membranes (Millipore). Membranes were then blocked in 5% Milk/ Tris-buffered saline + 0.01% Tween-20 (TBST), which was also used for subsequent incubation steps. Membranes were probed overnight at 4°C with antibodies against Polθ (rabbit polyclonal, raised against a GST-fusion protein encoding residues 1290-1389, Artios) and vinculin (mouse monoclonal, sc-73614, Santa Cruz). The membranes were then washed with TBST and incubated with HRP-conjugated anti-rabbit IgG secondary antibody (Thermofisher) and with IRDye 800CW Goat anti-mouse IgG Secondary antibody (Li-COR) for 1 h at RT in the dark. After washing twice in TBST, luminescent signals (from rabbit antibodies) were detected by ECL detection reagent (Thermofisher) and imaged on an Amersham Imager 600RGB; the fluorescent signals (from mouse antibodies) were detected using the Odyssey M Imager (Li-COR).

### Double thymidine (DT) block experiment

HeLa cells were plated as single cells for clonogenic survival assays, left to settle for 5 h and then incubated with 2 mM thymidine for 15 h, released for 9 h after a PBS wash and medium replacement, and incubated again with thymidine for 15 h. Then, cells were either released immediately after exposure to IR, or irradiated at 6 h after release. In all cases, either vehicle (DMSO) or ART558 was added 1 h before IR. To confirm the cell cycle position of the synchronized cells at the time of IR exposure, the same DT block protocol was applied in parallel to cells that were subsequently fixed in 70% ice-cold ethanol for DNA content analysis. Fixed cells were incubated with 50 μg/mL propidium iodide and 200 μg/mL RNase in PBS for 20 min at 37°C, and then analyzed using a Cytoflex (Beckman Coulter) cytometer. After exclusion of doublets, the distribution of cells according to their DNA content (PI intensity) was determined using FlowJo (BD) software.

### Nanoluciferase assay

The nanoluciferase MMEJ assay was carried out as described previously (22). Briefly, the Nano-luciferase MMEJ reporter construct (a linearized plasmid with protruding microhomology ssDNA ends engineered such to recombine into a functional luciferase construct only after correctly performed MMEJ) (22) was transfected into HEK293 cells together with a control firefly luciferase plasmid. Transfected cells were plated into 384-well plates already containing diluted compound and luminescence was read 24 h after transfection. Firefly and Nano-luciferase levels were detected using the Nano-Glo Dual-Luciferase Reporter Assay kit (Promega) and luminescence was measured with a Clariostar plate reader (BMG Labtech). The nanoluciferase signal was normalized to the firefly luciferase signal to correct for cell density and transfection efficiency.

### Immunofluorescence staining for DNA damage foci

Cells were seeded into 96-well thin bottom imaging plates (Perkin Elmer) and left to settle for a minimum of 6 hours at 37°C before treatment. ART558 was added 1 h prior to IR and cells were irradiated as described above. At specific time points after IR, cells were fixed in 4% paraformaldehyde for 15 minutes at room temperature. Cells were permeabilized in PBS with 1% BSA 0.5% Triton and 1% goat serum for at least 30 min. Cells were then incubated overnight at 4°C using any of the following primary antibodies diluted in PBS 1% BSA: rabbit anti RAD51 (Santa Cruz # sc-8349; 1/1500), rabbit anti-γH2AX (Novus Biologicals #NB100-2280; 1/2000), mouse anti-53BP1 (BD Biosciences # 612523: 1/1000) or mouse anti-dsDNA (to visualize micronuclei; SantaCruz #HYB331-01: 1:800). Following three PBS washes, cells were incubated for 90 min at room temperature with secondary antibodies diluted in PBS/1% BSA: Alexa488 goat anti-rabbit (Life Technologies # A11070; 1/1200) or Alexa594 goat anti-mouse (Life Technologies # A11020; 1/1200). DAPI (Sigma-Aldrich/Merck # D9542) was also added at 0.5 µg/mL final concentration at this point. Following three PBS washes, plates were imaged using an automated confocal microscope (GE Healthcare IN Cell 6000) and foci were counted using the Workstation software (GE Healthcare).

### Microsome assay

Metabolic stability of compounds in mouse and rat liver microsomes was conducted at WuXi AppTec Co. The compounds (ART558 and ART899) and controls (testosterone, diclofenac and propafenone) diluted in 100 mM potassium phosphate buffer (pH 7.4) were mixed with either mouse or rat microsome solution (Xenotech) at a 1:10 volume ratio (final concentration of compounds and microsomes, 1 µM and a 0.5 mg protein/mL, respectively), and incubated for 10 min at 37 °C. NADPH was then added to start the reactions. At different time intervals, ¾ of the final volume cold acetonitrile (containing 100 ng/mL tolbutamide and 100 ng/mL labetalol as internal standards) was added to stop the reaction. After centrifugation, the supernatants were analyzed by LC/MS/MS. A first order kinetics equation was used to calculate the half-life (t_1/2_), which was in turn used to calculate the intrinsic clearance based on the presence of the parent compound, according to the following equation: CLint = (0.693 / t_1/2_) x (1 / mg x mL^-1^ microsomal protein in reaction system) (μl x min^-1^ x mg^-1^), as previously described (24).

### In vivo studies

For the pharmacokinetics study, Nu/Nu mice (6-8 weeks) were dosed orally for 11 days with ART899 dissolved in 5% DMSO, 5% Ethanol, 20% TPGS, 30% PEG400, 40% water. Plasma samples were collected at various time points and ART899 levels were determined by CEMAS (Wokingham, UK) using standard protocols with a calibration curve and LC-MS/MS (API 5500) readout, and the free (non-protein bound) fraction of inhibitor in plasma was calculated. The pharmacokinetics study was carried out by WuXi AppTec (Shanghai, China).

The *in vivo* efficacy study was performed by Charles River (Morrisville, North Carolina). For the efficacy study, HCT116 cells (5 × 10^6^ in 100 µl PBS) were injected subcutaneously into female athymic nude mice (Crl:NU(NCr)-Foxn1nu, 8 – 12 weeks old). Fifteen days after tumor implantation –when average tumour size approached the target range of 80-180 mm^3^ – designated as day 1 of the study– tumour-size-matched animals (average tumour size 120.6-121.4 mm^3^) were sorted into the four treatment groups: Vehicle, ART899, 10 × 2 Gy + vehicle, and 10 × 2 Gy + ART899. Mice were treated with 150 mg/kg ART899 (dissolved as above) given by oral gavage BID for 12 days. Radiation was given on days 1-5 and 8-12 at 2 Gy per fraction targeted to the tumor while the mice were under isoflurane anesthesia. Tumour size and weight measurements were taken by one investigator not blinded to treatment group. Animals were weighed daily on days 1-5, then twice per week until the completion of the study. The endpoint for each group was a mean tumor size of 1500 mm^3^ or a maximum of 49 days. As such, the unirradiated treatment arms (vehicle or ART899 alone) were terminated at day 21, the 10 × 2 Gy arm on day 39, and the combination arm on day 49. The mice were observed frequently for overt signs of any adverse, treatment-related side effects, and clinical signs were recorded when observed. Any animal with weight loss exceeding 30% for one measurement, or exceeding 25% for three measurements, was euthanized as a treatment-related death. Acceptable toxicity was defined as a group mean body weight (BW) loss of less than 20% during the study and no more than 10% treatment-related death. Clinical observations, (including necropsies of main organs, especially spleen, liver and lymph nodes) were recorded at the time all available animals in each group were sampled. In the efficacy study, we did not observe any treatment-related death, nor significant decrease of body weight. In particular, a minimal reduction in body weight was associated to the IR treatment, and the group treated with IR + ART899 displayed a mean and distribution of body weight comparable to the group treated with IR alone. Necropsy analysis did not evidence any abnormality associated to the IR + ART899 combined treatment.

For the pharmacokinetics study, procedures related to animal handling, care and experimental treatments were performed according to the guidelines approved by the Institutional Animal Care and Use Committee (IACUC) of WuXi AppTec Co. following the guidance of the Association for Assessment and Accreditation of Laboratory Animal Care (AAALAC). The efficacy study performed by Charles River complied with the recommendations of the Guide for Care and Use of Laboratory Animals with respect to restraint, husbandry, surgical procedures, feed and fluid regulation, and veterinary care. The animal care and use program at Charles Rivers Discovery Services is accredited by AAALAC.

### Statistical analysis

Results are shown as the mean +/- standard deviation of a minimum of three observations unless otherwise indicated in the figure legends. Two-tailed t-tests were used to calculate statistical significance unless indicated otherwise; a p-value of < 0.05 was considered as statistically significant. The linear quadratic model, S = exp(αD – βD^2^), was used to fit clonogenic survival graphs, with S denoting survival, D the IR dose in Gy. Graphpad Prism was used for all statistical calculations and curve fitting.

## RESULTS

### ART558 induces Polθ-dependent radiosensitization in cancer cell lines

We have recently developed ART558, a new and specific Polθ inhibitor, which displayed synthetic lethality with BRCA loss (22). In the present study, we investigated whether Polθ inhibition could serve as an anti-cancer therapeutic agent beyond the context of BRCA deficiency, in combination with radiotherapy. We first assessed the effect of ART558, alone and in combination with ionizing radiation (IR), in three tumor cell lines that do not harbor mutations in BRCA-1 or BRCA-2 genes (25): HCT116 (colorectal), H460 (lung) and T24 (bladder). Using colony formation assays we confirmed that ART558 does not cause any effect in the absence of ionizing radiation (IR) (Figure 1A), but effectively radiosensitizes all three BRCA-proficient cell lines from 1 µM concentration (Figure 1B; SER_10_ (sensitization enhancement ratio at 10% survival) > 1.2 at 1 µM ART558 in all three cell lines).

**Figure 1.**
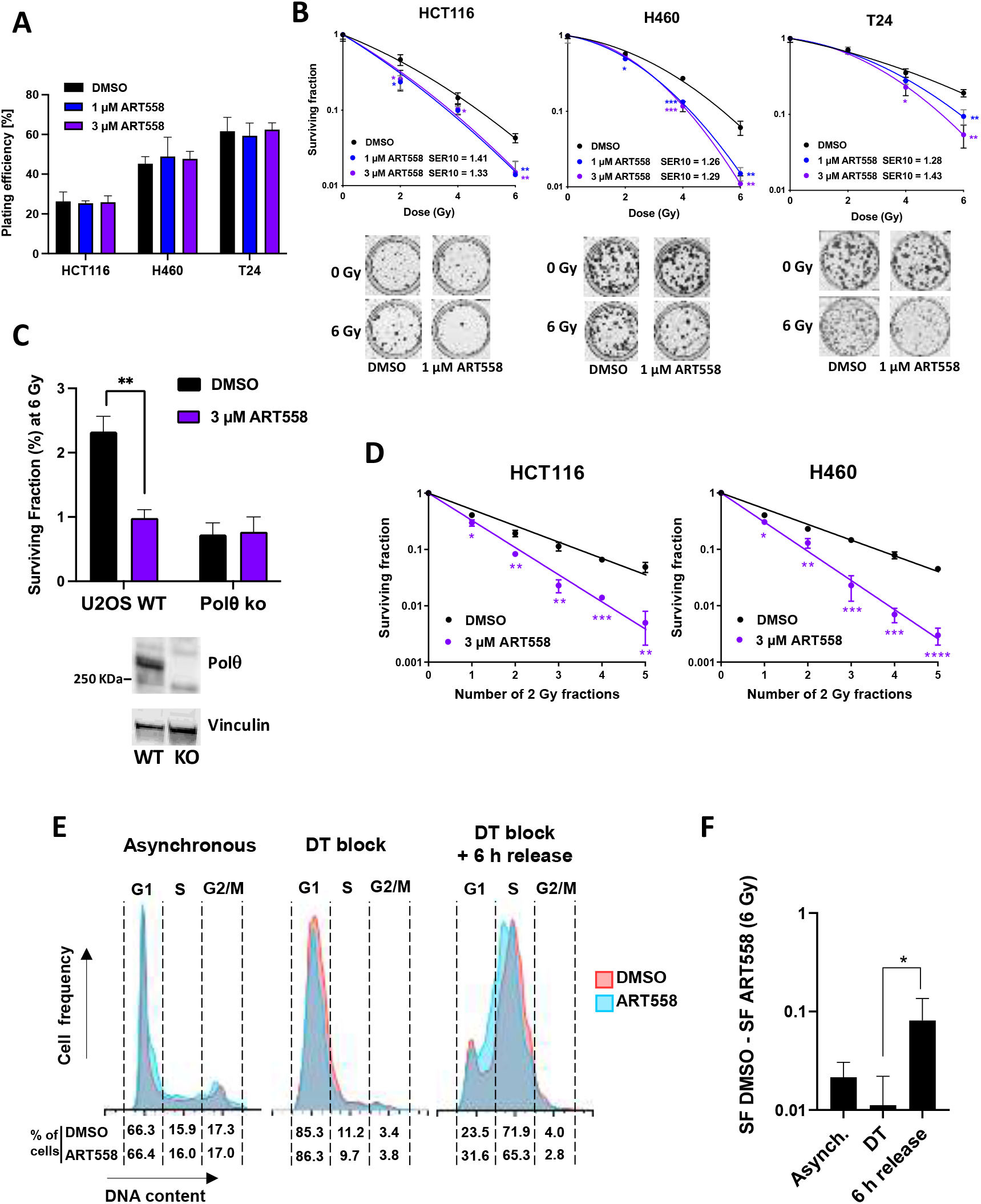
The Polθ inhibitor ART558 radiosensitizes tumor cells. (**A-B**) Clonogenic survival of HCT116, H460 and T24 cells treated with ART558 and/or IR. (**A**) Plating efficiency for unirradiated cells. (**B**) Surviving fractions as a function of the irradiation dose. SER10: Sensitization Enhancement Ratios for 10% survival. Representative wells for 0 Gy and 6 Gy +/- 1 µM ART558 are shown for each cell line. (**C**) Clonogenic survival of U2OS WT and Polθ KO cells treated with 3 µM ART558 and 6 Gy IR. Bar graphs show the surviving fraction at 6 Gy. The Western blot insets confirm the lack of Polθ expression in the U2OS Polθ KO cells. (**D**) Clonogenic survival of H460 and HCT116 treated with 3 µM ART558 and 2 × 5 Gy (2 Gy once per day for 1 to 5 days). (**E-F**) Clonogenic survival following irradiation and treatment with 3 µM ART558 of synchronized HeLa cells. (**E**) Representative histograms showing the cell cycle distribution at the time of IR (synchronized in G1 by DT block or after 6 h release from DT block, compared to asynchronous cultures). (**F**) Degree of radiosensitization estimated by the difference between the surviving fraction of DMSO- and ART558-treated cells after IR (SF DMSO – SF ART558). All data correspond to average +/- SD from triplicate wells (representative from 3 separate experiments); (* p<0.05; ** p<0.01; *** p<0.001; **** p<0.0001).

Because 3 µM gave a comparable or stronger response than 1 µM whilst not affecting viability without radiation, this ART558 concentration was chosen for subsequent experiments. U2OS Polθ CRISPR knockout (KO) cells were more sensitive to IR than parental wild-type (WT) U2OS cells and were not radiosensitized by ART558, whereas the WT U2OS cells were significantly radiosensitized, confirming that the radiosensitizing effect of ART558 is Polθ-specific (Figure 1C). Since radiotherapy is given over multiple fractions in clinical settings (26), we tested ART558 combined with fractionated radiotherapy, and showed that the radiosensitizing effect of ART558 increases with the number of fractions (Figure 1D). For H460 cells there was an up to a 14-fold decrease in survival with 5 × 2 Gy in the presence of ART558 compared with IR alone. For HCT116 cells, this factor was up to 10-fold.

As MMEJ functions mainly during the S and G2 cell cycle phases (27), we hypothesized that Polθ inhibition would have a greater effect in these cell cycle phases. To investigate this, we used the cervical cancer cell line HeLa due to its suitability for cell cycle synchronization using double thymidine (DT) block. As expected, we found that HeLa cells irradiated when they are mostly traversing S phase are radiosensitized by ART558 to a higher extent than cells irradiated whilst synchronized in G1 (Figure 1E, F). Since fractionated radiotherapy allows for cells to move to different cell cycle phases where they might be more sensitive to Polθ inhibition (26), this finding may partly explain our observation that ART558-mediated radiopotentiation is higher with multiple IR fractions.

### Increased DNA damage after ART558 and radiation

To clarify whether the radiosensitizing effect of ART558 is exerted through an impairment in DSB repair, we assessed changes in both phosphorylated H2AX (γH2AX) and the NHEJ factor 53BP1. In the presence of ART558, we observed an increase in both γH2AX and 53BP1 foci 24 hours after IR (Figure 2A, B), demonstrating that ART558-mediated inhibition of Polθ results in a higher number of persistent IR-induced DSBs. Polθ has been shown to directly antagonize the binding of the HR repair factor RAD51 to ssDNA (7) and, accordingly, Polθ depletion has previously been shown to increase the formation of RAD51 foci in HR-deficient cells (28,29). To determine whether Polθ inhibition affects the recruitment of RAD51 to IR-induced DSBs in HR-proficient cells, we quantified the number of RAD51 foci at 6 hours after IR –when RAD51 foci number are at their peak (30)– but we did not observe any significant difference (Figure 2C, D).

**Figure 2.**
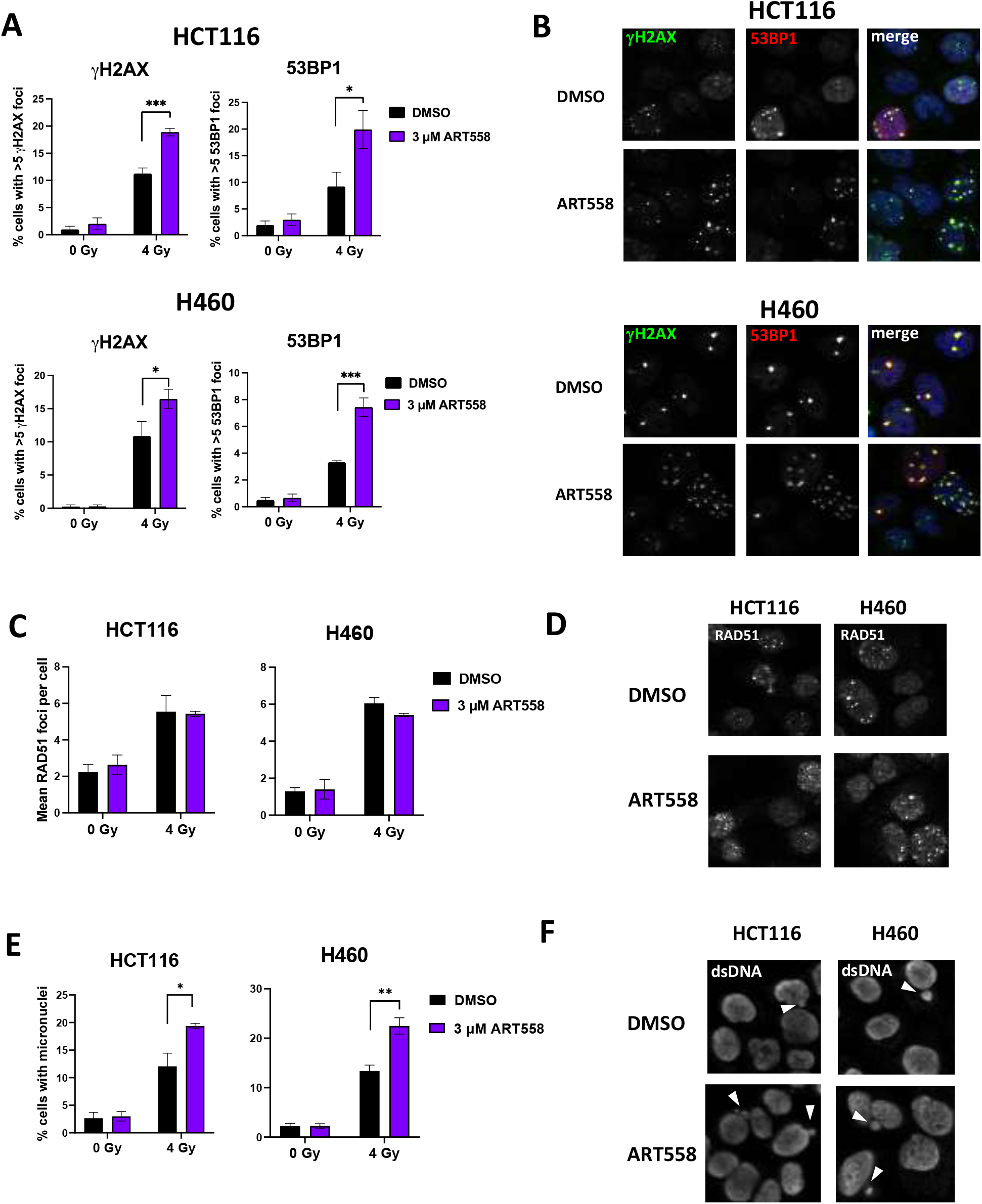
ART558 treatment leads to increased residual IR-induced DNA damage foci. γH2AX and 53BP1 foci in H460 and HCT116 treated with 3 µM ART558 and 4 Gy IR, assessed 24 h after IR. γH2AX: green; 53BP1: red. (**B**) Representative images from irradiated cells from the experiment described in (A). (**C**) RAD51 foci in H460 and HCT166 treated with 3 µM ART558 and 4 Gy IR, assessed 6 h after IR. (**D**) Representative images from irradiated cells from the experiment described in (C). (**E**) Micronuclei in H460 and HCT116 cells treated with 3 µM ART558 and 4 Gy IR, assessed 48 h after IR. (**F**) Representative images from irradiated cells from the experiment described in (E). Micronuclei are indicated with arrow heads. Data points indicate the mean +/- SD from triplicate wells and graphs are representative from 3 separate experiments.

### Effect of ART558 under hypoxic conditions

Hypoxic cells can be over three times more radioresistant than normoxic cells (31,32) and for this reason tumor hypoxia is an important mechanism of radiotherapy resistance. Therefore, we tested ART558 combined with IR at two low oxygen concentrations: 0.5% and <0.1% oxygen. The results confirmed that hypoxia induces significant resistance to IR (oxygen enhancements ratios (OERs) of 1.31 and 1.54 for 0.5% and < 0.1% oxygen, respectively: Suppl. Figure 1). Nonetheless, we found that even under severe hypoxia (<0.1% oxygen), ART558 confers significant radiosensitization, with SER_10_ values just slightly lower than those in normoxic conditions (Figure 3).

**Figure 3.**
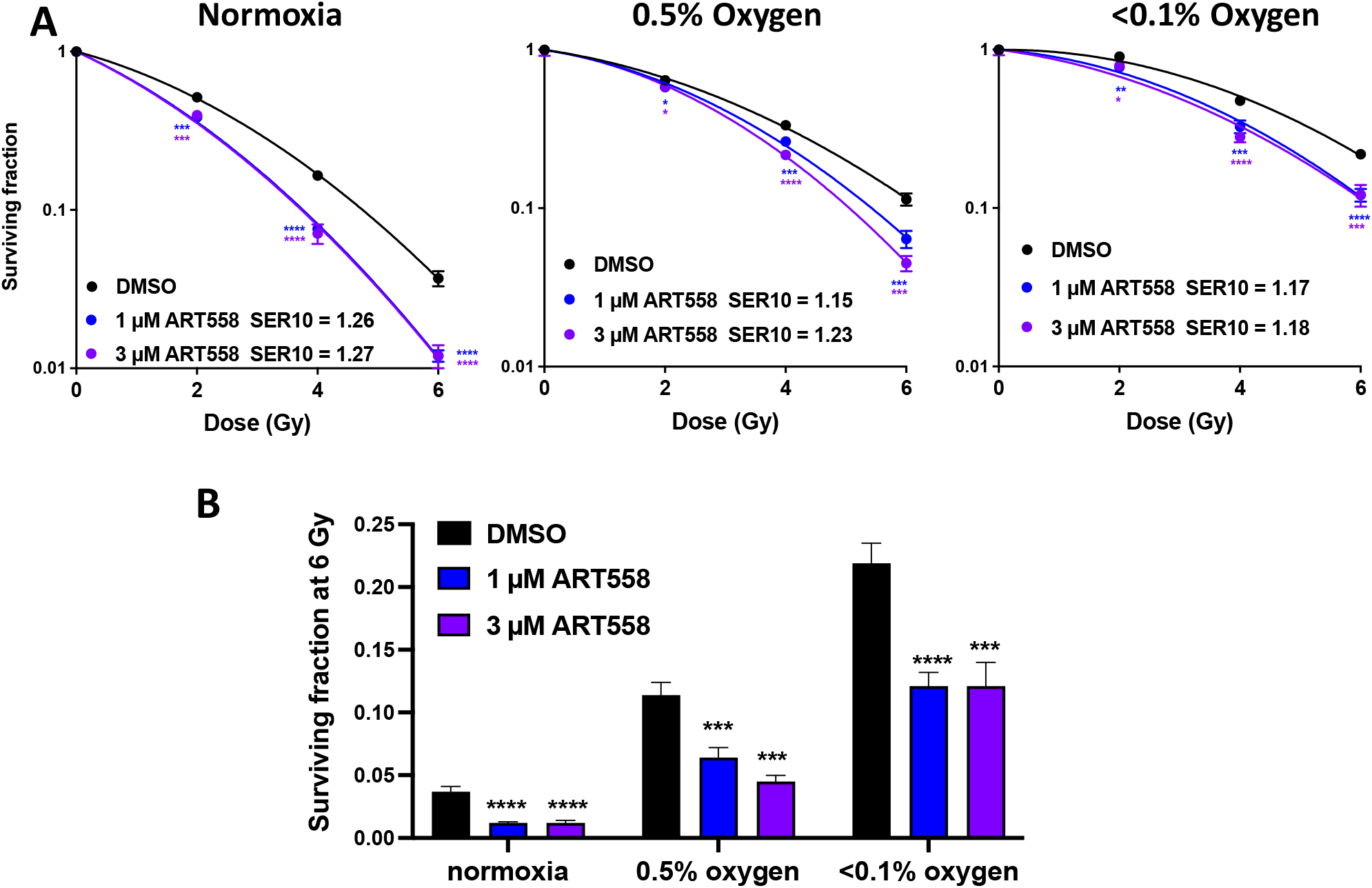
ART558-mediated radiosensitization under hypoxic conditions. (**A**) Clonogenic survival of H460 cells irradiated upon hypoxia (0.5% and <0.1% oxygen). For oxygen enhancement ratios see Suppl. Figure 1. The 6 Gy data points are replotted in a bar graph in panel (**B**) to allow better visual comparison between the different treatment arms. * p<0.05; ** p<0.01; *** p<0.001; **** p<0.0001.

### Polθ inhibition combined with radiation is well tolerated and leads to reduced tumor growth *in vivo*

Next, we went on to determine the efficacy of Polθ inhibition in combination with radiation *in vivo*. As ART558 has been previously shown to be unsuitable for *in vivo* treatment due to poor metabolic stability (22), we utilized an optimized derivative, ART899 (Fig 4A). ART899 is a deuterated form of ART812, the ART558 derivate shown to have efficacy in BRCA1/SHLD2-deficient tumor xenograft studies (22). Clearance values obtained using microsome stability assays showed that ART899 has a greatly improved metabolic stability compared to ART558 (Fig 4A). To confirm that ART899 is an effective and specific Polθ inhibitor, we tested its activity in *in vitro* assays. First, we measured the inhibition of cellular MMEJ activity in a luciferase assay previously used to demonstrate ART558 activity (22). This showed that ART899 has a cellular IC50 of approximately 180 nM (Figure 4B-C), which is comparable to that of ART558 (IC50 = 150 nM) (22).

**Figure 4.**
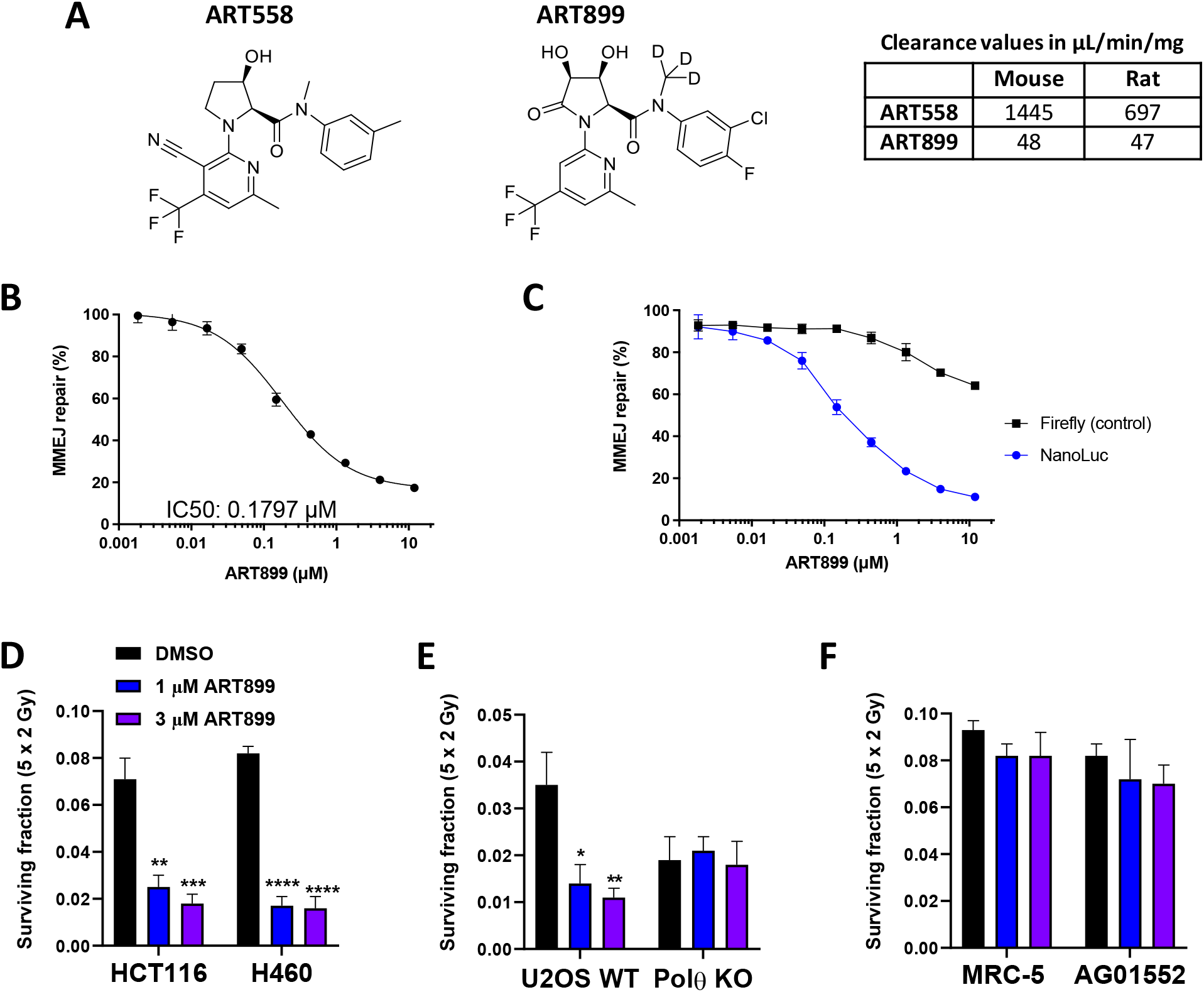
Characterization of ART899 as a specific and potent Polθ inhibitor with improved stability. (**A**) Chemical structures of the Polθ inhibitors ART558 and ART899. The table shows the *in vitro* intrinsic clearance values of ART558 and ART899 after exposure to rat and mouse liver microsomes. (**B**) Nano-luciferase MMEJ assay showing ART899-mediated inhibition of MMEJ activity. The nano-luciferase readings were normalized to control luciferase (firefly) readings, and these were then normalized to DMSO. Data points show the mean +/- SEM of two independent experiments. Data points show the mean +/- SEM of two independent experiments. (**C**) Confirmation of MMEJ assay specificity. Same experiment described in (B) but showing both the nanoluc and firefly readings normalized to their own DMSO reading, confirming negligible inhibition by ART899 of the control firefly luciferase signal. (**D**) Clonogenic survival of H460 and HCT116 cells treated with ART899 and irradiated with 2 × 5 Gy. Graphs show the surviving fraction at 5 × 2 Gy. (**E**) Confirmation of ART899 specificity in U2OS WT and Polθ KO cells. Cells were treated as described in (D). (**F**) Effect of ART899 in non-cancerous cells. MRC-5 and AG01552 fibroblasts were treated as described in (D). The effect of ART899 in unirradiated cells from (D-F) is shown in Suppl. Figure 2. Graphs shown in (D-F) correspond to average +/- SD from triplicate wells (representative from 3 separate experiments). * p<0.05; ** p<0.01; *** p<0.001; **** p<0.0001.

Since our earlier results had shown that ART558 is more effective when combined with multiple IR doses rather than a single IR dose (Figure 1), we investigated the radiosensitizing efficacy of ART899 in combination with fractionated IR (5 × 2 Gy). This showed that ART899 effectively radiosensitizes HCT116 and H460 cells at both 1 µM and 3 µM (Figure 4D). At the highest ART899 concentration (3 µM), the decrease in survival in H460 cells was 5-fold compared to IR alone; for HCT116 this factor was 4-fold. Similar to ART558, we did not observe any effect of ART899 in the absence of irradiation (Suppl. Figure 2).

Specificity for Polθ was further confirmed by showing a lack of radiosensitization in U2OS Polθ KO cells (Figure 4E). Prior to starting *in vivo* studies, it was important to confirm that ART899 does not radiosensitize non-cancerous cells. To this end, we tested ART899 combined with 5 × 2 Gy IR in two human fibroblast lines, MRC-5 and AG01552. As shown in Figure 4F, there was no radiosensitization in either of these cell lines, demonstrating the tumor specificity of this Polθ inhibitor.

We then confirmed that ART899 possesses a good *in vivo* pharmacokinetic profile, with a plasma concentration ≥ 1µM for several hours after oral administration in mice (Figure 5A). Finally, we tested ART899 in mice bearing HCT116 subcutaneous xenografts. The combination of ART899 treatment with fractionated irradiation (10 × 2 Gy) significantly improved tumor growth delay compared to radiation alone (Figure 5B-D and Suppl. Figure 3). Importantly, the combination treatment was well tolerated with mice displaying no weight loss, or other signs of distress or treatment-related adversity, compared to IR alone, during the course of the experiment and as per necropsy at the experiment end (Figure 5E and Suppl. Figure 3). Together, these data show that these novel Polθ inhibitors are effective tumor-selective radiosensitizers in both *in vitro* and *in vivo* settings.

**Figure 5.**
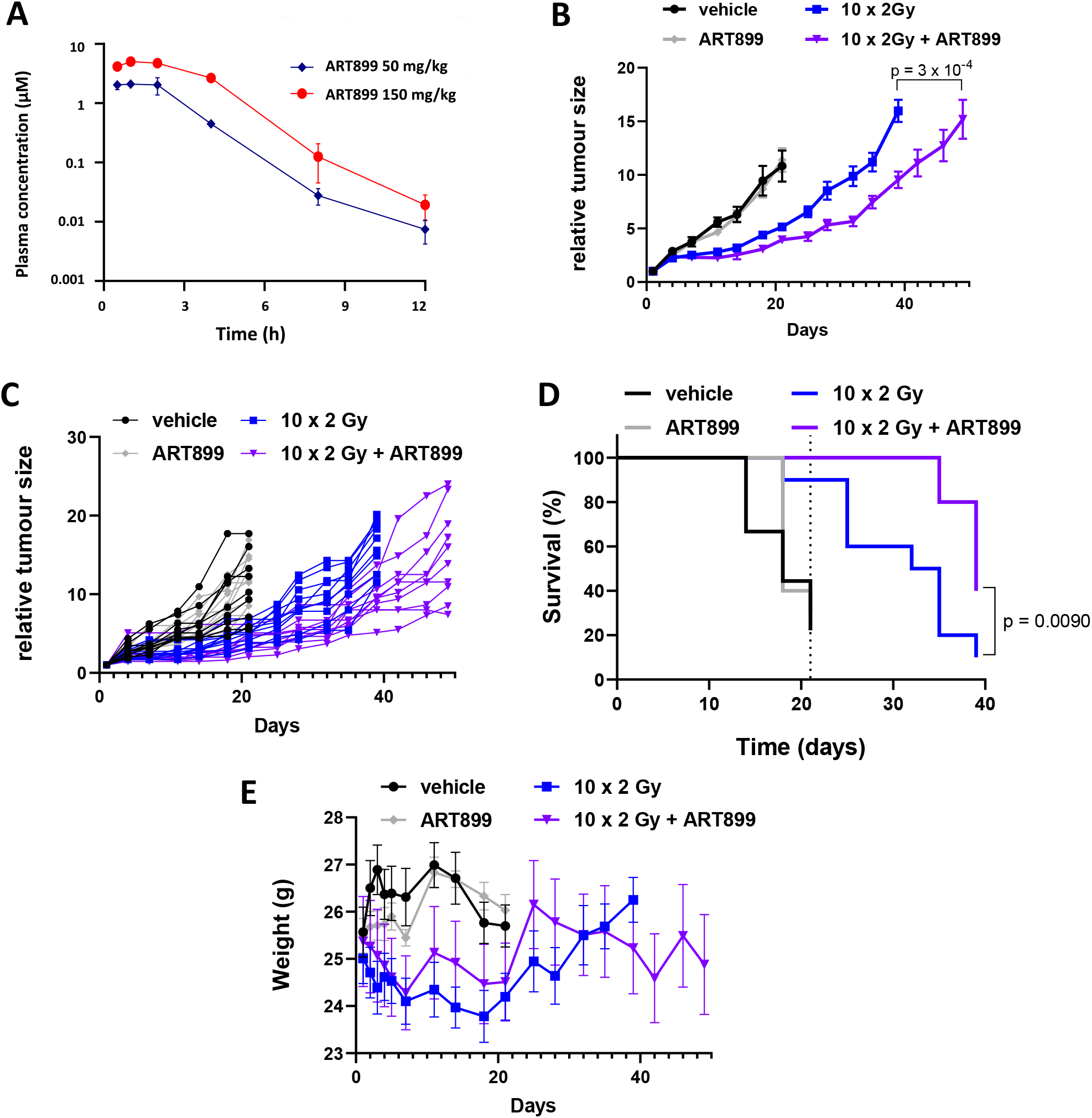
Polθ inhibitor ART899 combined with radiation causes significant tumor growth delay *in vivo* and is well tolerated. (**A**) ART899 plasma concentration following oral dosage of ART899 at 50 or 150 mg/kg. Mouse plasma samples (n = 3 per treatment group) were collected at 30 min, 1 h, 2h, 4h, 8h and 12h post last dose. (**B-E)** HCT116 tumor-bearing mice treated with 150 mg/kg Polθ inhibitor ART899 BID for 12 days and/or 10 × 2 Gy (days 1-5 & 8-12). Vehicle (n=9); ART899 (n=10); 10 × 2 Gy + vehicle (n=10); 10 × 2 Gy + ART899 (n=10). (**B**) Mean +/- SEM tumor size. p-value from mixed effect model and Dunnett post-test. Individual mouse graphs and comparison of tumor size at the latest common endpoint for 10 × 2 Gy *vs*. 10 × 2 Gy + ART899 are shown in Suppl. Figure 3. (**C**) Individual mouse graphs. (**D**) Kaplan Meier plot for a tumor size threshold of 1000 mm^3^. p-value from the log-rank (Mantel-Cox) test comparing IR alone and IR+ART558 is indicated. (**E**) Average mouse weight +/- SD from all treatment groups over time. Individual mouse weights are shown in Suppl. Figure 3.

## DISCUSSION

Here, we demonstrate the radiosensitizing effect of novel Polθ inhibitors ART558 and ART899. ART558, was previously shown to have a potent stand-alone anti-tumor effect in cancer cells with defects in HR and the Shieldin complex, and in combination treatment with PARP inhibitors (22). We now show that ART558 is also able to significantly radiosentize HR-proficient cells in a Polθ-specific fashion. In addition, we report for the first time ART899, a novel derivative of ART558 with increased stability and efficacy against Polθ *in vivo*. We confirmed that ART899 specifically inhibits Polθ MMEJ activity and that specifically radiosensitizes tumor cells with no effect on non-cancerous cells, consistent with the low or absent expression of Polθ in normal tissues and overexpression in many cancer cells (4,5,21).

We also report here a higher proportion of residual IR-induced γH2AX and 53BP1 foci upon pharmacological inhibition of Polθ linked to increased formation of IR-induced micronuclei, which strongly suggests that the mechanism whereby our Polθ inhibitors radiosensitize cancer cells is by an impairment in DSB repair which leads to lethal chromosomal rearrangements (33). Our finding that Polθ inhibition does not lead to a higher number of RAD51 foci is perhaps in contrast with earlier findings that showed an increase in RAD51 foci in HR-proficient, Polθ-depleted cells following radiation (7,20). These discrepancies may be explained by the fact that those experiments used RNA interference or genetic depletion rather than pharmacological inhibition. One plausible explanation is that, unlike Polθ depletion, Polθ inhibition preserves the integrity of the Polθ molecule, which would remain bound to the resected DSB and therefore would still be able to compete with RAD51 for its binding to DNA (7).

Importantly, we show that Polθ inhibition potently radiosensitizes cancer cells exposed to multiple fractions of radiation, both *in vitro* and *in vivo*. The translational relevance of this is that, in a clinical setting, radiotherapy is commonly administered in multiple doses –i.e., fractionated radiotherapy– which minimizes toxicity to normal tissues. In line with the known cell cycle selectivity of the MMEJ repair pathway (27), we confirmed that cells traversing S phase are more sensitive to Polθ inhibition than cells in G1 phase. Since cancer cells are known to transit through the cell cycle during the time between IR fractions –which increases the likelihood of cancer cells being irradiated at more radiosensitive cell cycle phases (26)– this finding provides mechanistic evidence supporting the advantage of delivering a fractionated radiotherapy regime when using Polθ inhibitors. In addition, we show for the first time that Polθ inhibition can radiosensitize tumor cells under hypoxia, a common feature of solid tumors that confers radioresistance (32). Our results therefore position Polθ inhibition as a promising strategy to improve the efficacy of radiotherapy regardless of the presence of tumor hypoxia.

Finally, we show here that combining ART899 with fractionated IR in xenograft models results in significantly improved tumor growth delay and is well tolerated. Polθ inhibition is being currently tested in a first-in-human clinical trial (NCT04991480) which is evaluating the safety and activity of Polθ inhibition in patients with solid tumors. This clinical trial could open new clinical avenues to explore the therapeutic modality we have discovered in this work. Our study broadens the potential therapeutic application of Polθ inhibition beyond its use as a therapy for tumors with defects in HR repair, with the promise to improve the efficacy of radiotherapy.

