## Supplementary material for "Small-molecule Polθ inhibitors provide safe and effective tumor radiosensitization in preclinical models": Suppl. Figures

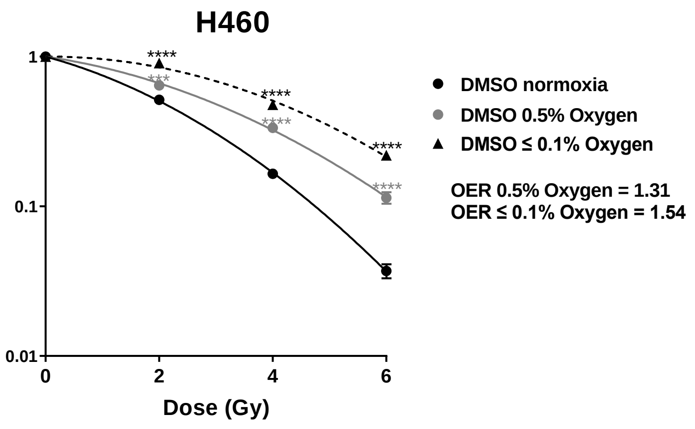


**Suppl. Figure 1.** Accompanies Figure 3 (Confirmation of resistance to IR induced by hypoxia). Clonogenic survival graph comparing control (DMSO) curves from Figure 3A. OER: oxygen enhancement ratios.*** $p<0.001$; **** $p<0.0001$.

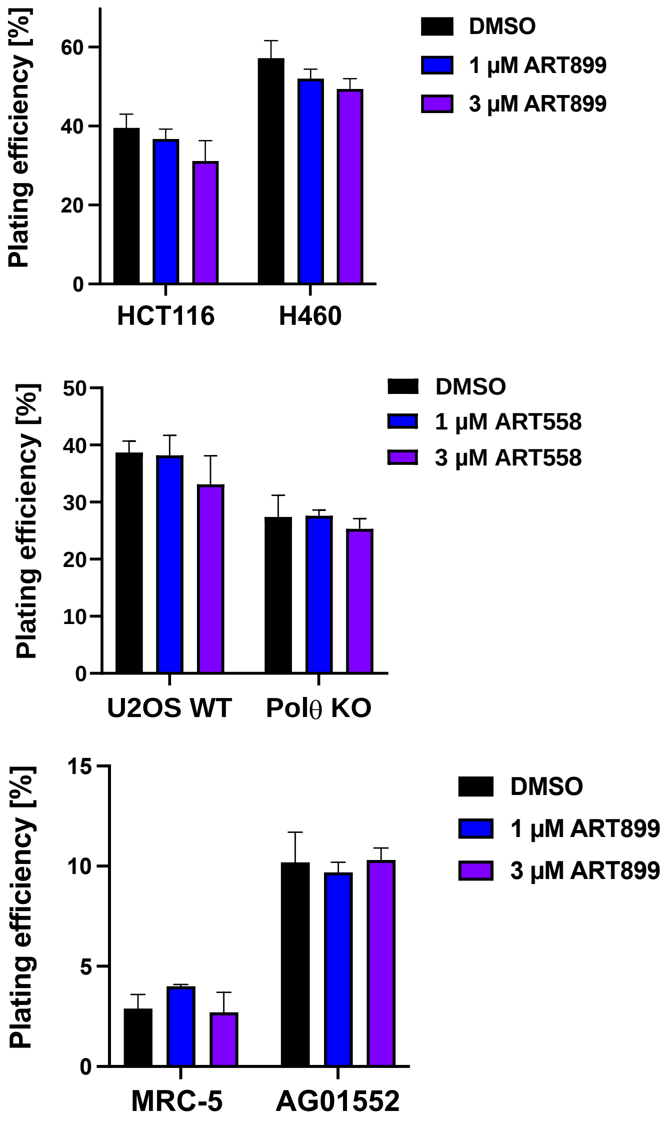


**Suppl. Figure 2.** Accompanies Figure 4 (Characterization of ART899 as a specific and potent Polθ inhibitor with improved stability). Effect of ART899 in unirradiated cells from experiments described in Figure 4D-F.

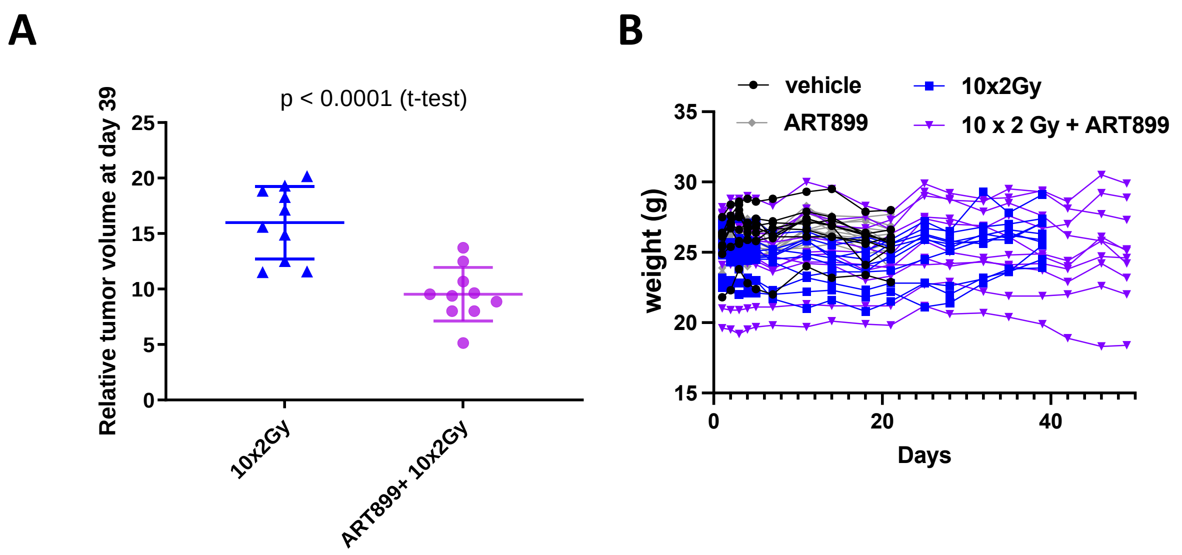


**Suppl. Figure 3.** Accompanies Figure 5 (ART899 combined with radiation causes significant tumor growth delay in vivo and is well tolerated).

**(A)** Comparison of tumor size at day 39, from Figure 5B.

**(B)** Individual mouse graphs from Figure 5E.
